# Robust coordination of collective oscillatory signaling requires single-cell excitability and fold-change detection

**DOI:** 10.1101/2021.09.02.457527

**Authors:** Chuqiao Huyan, Alexander Golden, Xinwen Zhu, Pankaj Mehta, Allyson E. Sgro

## Abstract

Complex multicellular behaviors are coordinated at the level of biochemical signaling networks, yet how this decentralized mechanism enables robust control in variable environments and over many orders of magnitude of spatiotemporal scales remains an open question. A stunning example of these behaviors is found in the microbe *Dictyostelium discoideum*, which uses the small molecule cyclic AMP (cAMP) to drive the propagation of collective signaling oscillations leading to multicellular development. The critical design features of the *Dictyostelium* signaling network remain unclear despite decades of mathematical modeling and experimental interrogation because each model makes different assumptions about the network architecture and in general, normalizing models for direct comparison presents a major challenge. We overcome this challenge by using recent experimental data to normalize the time and response scales of five major signal relay network models to one another and assess their ability to recapitulate experimentally-observed population and single-cell dynamics. We find that to successfully reproduce the full range of observed dynamical behaviors, single cells must be excitable and respond to the relative fold-change of environmental signals. This suggests these features represent robust principles for coordinating cellular populations through oscillatory signaling and that single-cell excitable dynamics are a generalizable route for controlling population behaviors.

## INTRODUCTION

Cellular populations often work together to engage in complex emergent multicellular behaviors. These behaviors range from the computations performed by populations of neurons [1], to coordinated growth within a biofilm composed of thousands of bacteria [2, 3]. Remarkably, these population-level behaviors are often coordinated by biochemical intracellular signaling networks. This decentralized control of population-wide behaviors by internal signaling networks is impressive given that individual cells often have access to very limited information about their immediate surroundings. Furthermore, cell-cell communication mechanisms commonly utilize only a handful of signaling molecules, limiting information transmission between cells. This raises natural questions about the strategies cells employ to use this limited information to robustly control emergent population behaviors in a wide variety of contexts.

To address this question, we focus on one of the most notable and extensively-studied examples of collective multicellular phenomena, the starvation response of the social amoeba *Dictyostelium discoideum*, where cells transition from a free-living unicellular state to a multicellular aggregate [4, 5]. Upon starvation, *Dictyostelium* cells initiate a developmental program where some cells start releasing pulses of cyclic AMP (cAMP) into the external space [6–8]. These oscillatory pulses are relayed through the population, creating external cAMP waves that act as a chemoattractant directing cell migration towards the source of cAMP during aggregation. These waves allow *Dictyostelium* to self-organize and develop into a stalk-and-spore structure in complex and ill-defined environments over group sizes that can vary by many orders of magnitude [4]. Identifying how this level of control is achieved is still an open question as most studies on design principles that coordinate multicellular phenomena have focused on developmental networks in metazoans that use well-defined morphogen gradients to pattern their development [9–13].

Even in a well-studied model organism such as *Dictyostelium*, identifying the critical features of single-cell signaling networks that drive emergent population-level behaviors poses a tremendous challenge as it requires identifying the underlying network architecture. To address this challenge, there has been a grand tradition of mathematical modeling to link the observed molecular and cellular behaviors to specific network architectures that can drive group-wide phenomena. These models originally focused on population-wide oscillations based on biochemical data [14–16]. Recent experimental advances exploiting genetically-encoded cAMP sensors and microfluidic platforms have enabled measurements of single-cell signaling dynamics in both isolated cells and collectively-behaving populations, helping shed light on how the single-cell signaling network may be configured. [7, 8, 17, 18].

While all the models describing *Dictyostelium* signaling reproduce population-wide oscillations, not all of them produce the more complex single-cell and population behaviors revealed by these recent experiments. Furthermore, they make conflicting assumptions about the architectural features of single-cell signaling networks that lead to different behaviors beyond coordinated oscillatory signaling. Resolving these issues represents a major conceptual challenge because these models have different time and response scales with different arbitrary units. Here, we address this challenge by exploiting recent single-cell measurements: when individual *Dictyostelium* cells are exposed to a threshold level of external cAMP stimulation, they respond with a characteristic adaptive pulse of internal cAMP with a reproducible timescale and magnitude. We show below that we can use this characteristic cellular response to normalize the time and response amplitude scales of different mathematical models to one another to directly compare their behaviors. This allows us to screen for how accurately these mathematical models recapitulate experimentally-observed population and single-cell signaling phenomena, and to identify the key signaling network features and single cell properties that are critical for driving population behaviors. We find that single-cell excitability and fold-change detection are critical singlecell properties for robustly coordinating population-wide oscillations that can be modulated through an external medium.

## I. RESULTS

### A. Simulation framework for comparing mathematical models of *Dictyostelium* signal relay

We analyze five major models that describe the *Dictyostelium* signal relay network (Figure 1). These models were developed over the last thirty-five years based on experimental observations and include both bottom-up mechanistic models that seek to directly relate variables to specific proteins [14, 16] and top-down phenomenological models that seek to abstract away molecular details yet still capture the observed dynamical behaviors [7, 8, 17]. In order to identify common features required to reproduce the observed dynamical behaviors and illuminate the design principles required for collective coordination, we classified models according to architecture and developed a unified simulation framework that allowed us to compare their behavior to both population and single-cell level experimental data. The different signaling network architectures are built up from different network design features. These features include control loops, such as positive feedback, negative feedback, and incoherent feedforward loops that can be integrated with one another. Other design features are logarithmic sensing receptors that sense order of magnitude changes in external cAMP inputs, and treating each cell as a phase oscillator. Furthermore, given recent work suggesting that noise plays a prominent role in population-wide coordination [7, 8], we constructed stochastic generalizations of the original deterministic models to understand the effect of noise on population-level behaviors. The models are implemented in Python using standard methods for solving ordinary differential equations (Figure 1) and their stochastic counterparts. The models are summarized in detail in the *Supplementary Materials*, highlighting the underlying assumptions the models make about the network architectures.

**FIG. 1:**
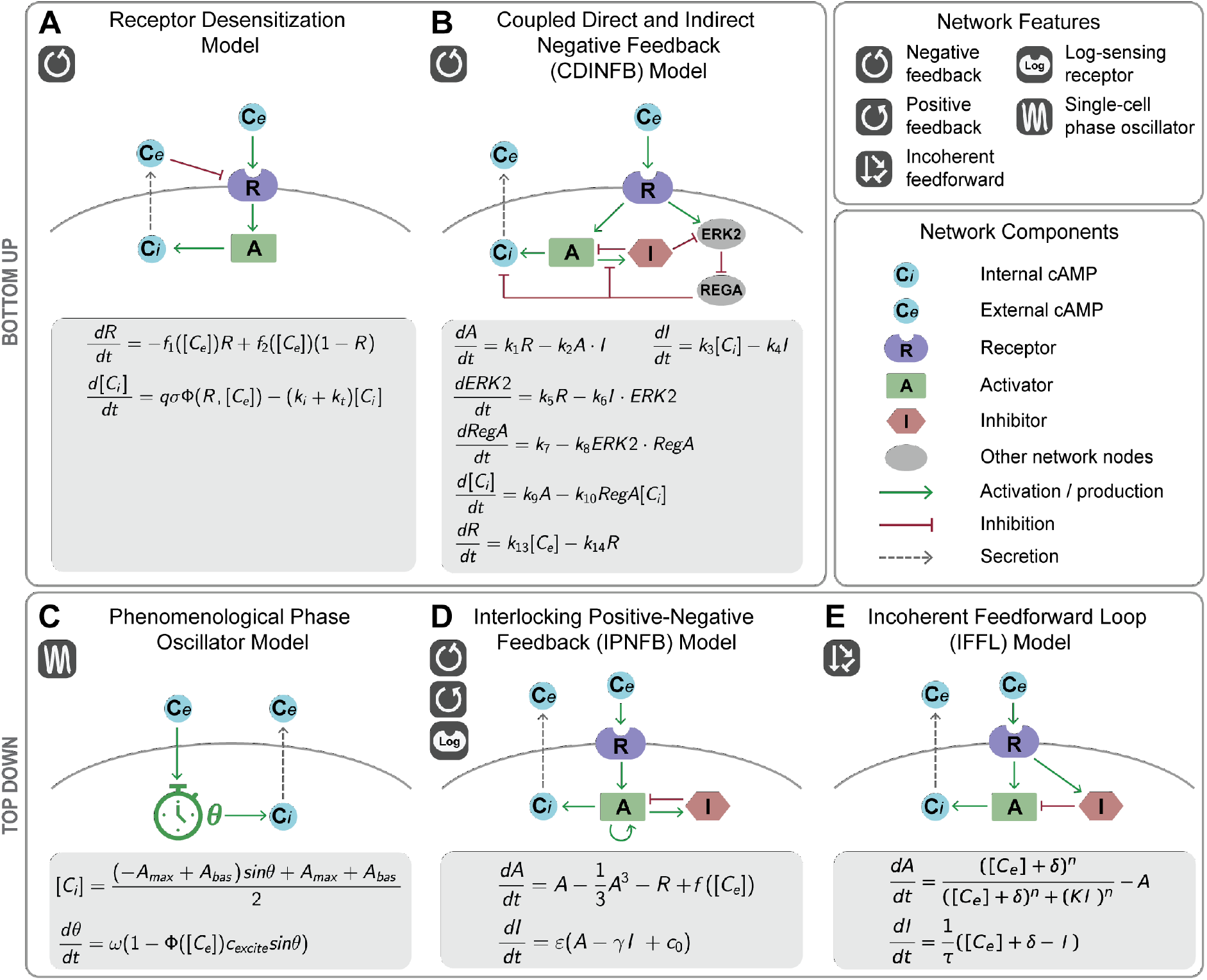
Five major mathematical models describing dynamical signaling relay behaviors in *Dictyostelium*. Models are abstracted as network diagrams with common signaling network components. The activator in each model either stands for the enzyme that directly produces cAMP (adenylyl cyclase) or internal cAMP itself. The top panel illustrates the bottom-up models based on receptor desensitization **(A)** and a coupled direct and indirect negative feedback (CDINFB) architecture **(B)** that use kinetic equations based on biological signaling networks. The bottom panel shows top-down models that use well-studied mathematical equations to recapitulate dynamics from recent experiments: a coupled phase oscillator model **(C)**, an interlocking positive-negative feedback (IPNFB) model that adapts the FitzHugh-Nagumo model framework and integrates a logarithmic sensing receptor module **(D)**, and an incoherent feedforward loop (IFFL) model **(E)**.

To permit direct comparison between the models, as well as to experimental data, we needed a common scale for three key parameters: external cAMP level, internal cAMP level, and time. To accomplish this, we normalized each model to match a characteristic experimental behavior: in response to a low-level 1 nM step of cAMP, single cells produce a spike of internal cAMP with reproducible height and width before returning to baseline (Figure 2A) [8]. For each model, we set 1 external arbitrary unit of cAMP equal to either the amount designated as 1 unit or 1 nM in the model [7, 8, 17] or the minimum amount of [*cAMP*]_*e*_ that produced a robust spike of internal cAMP [14, 16] (Figure 2B-F). All models except the Phase Oscillator model display this spike, whereas phase oscillators are designed to bifurcate to oscillations at high levels of external stimulation, so we used either this spike or the oscillations to establish common internal cAMP levels and time units. Specifically, we set the internal cAMP response level of one arbitrary unit to the height of the Phase Oscillator oscillations at the equivalent of 10 μm external cAMP (Figure 2B) or the height of the internal cAMP response to 1 external arbitrary unit of cAMP (Figure 2C-F). Similarly, we set one arbitrary time unit to the intrinsic oscillation timescale in the Phase Oscillator model (Figure 2B) [7], the time to return to 5% of the internal cAMP amplitude over the post-stimulation lowest level of internal cAMP in the Receptor Desensitization, the CDINFB, and the IFFL models (Figure 2C-E) [14, 16, 17], and the adaptive spike timescale parameter set in the original paper in the IPNFB model (Figure 2F) [8].

**FIG. 2:**
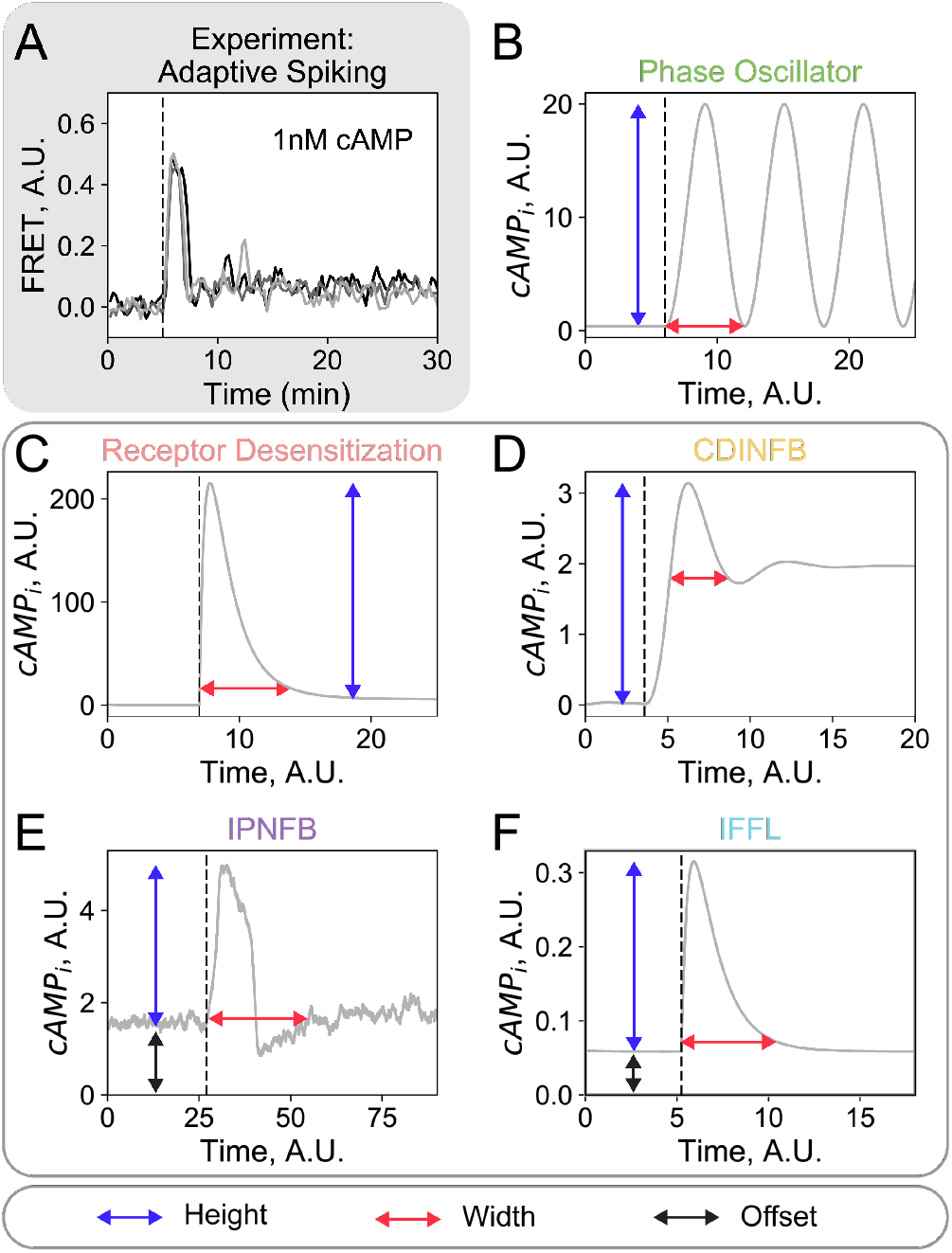
Normalizing model response and timescales to the characteristic adaptive spike in single cells. **(A)** Experimental data shows single *Dictyostelium* cells display an adaptive internal cAMP spike in response to a 1 nM external cAMP step input (n = 3 example cells). Data adapted from Sgro, et al. [8]. In each model, the time and response amplitude normalization parameters (time: red arrows; amplitude: blue arrows) are set to the height and width of either the oscillations **(B)** or the spike the model displays in response to 1 external arbitrary unit of cAMP **(C – F)**. In the IPNFB and IFFL models the response amplitude is offset to normalize their basal internal cAMP levels to 0 (offset: black arrows) **(E and F)**. In both the experimental data and model results, external cAMP is applied at the black dashed line. Input external cAMP concentrations were determined as described in the main text. Units shown in **B-F** are the original model units.

### B. Comparison of mathematical models to population and single cell experimental data

We compared dynamical behaviors displayed by the five different models against nine key dynamical behaviors observed in cellular populations and isolated single cells. The performance of the five models is summarized in Table I and full model details and simulation results can be found in the *Supplementary Materials*. For brevity, in the main text we limit our discussion to the four most informative behaviors for distinguishing between different network design features.

**TABLE I:**
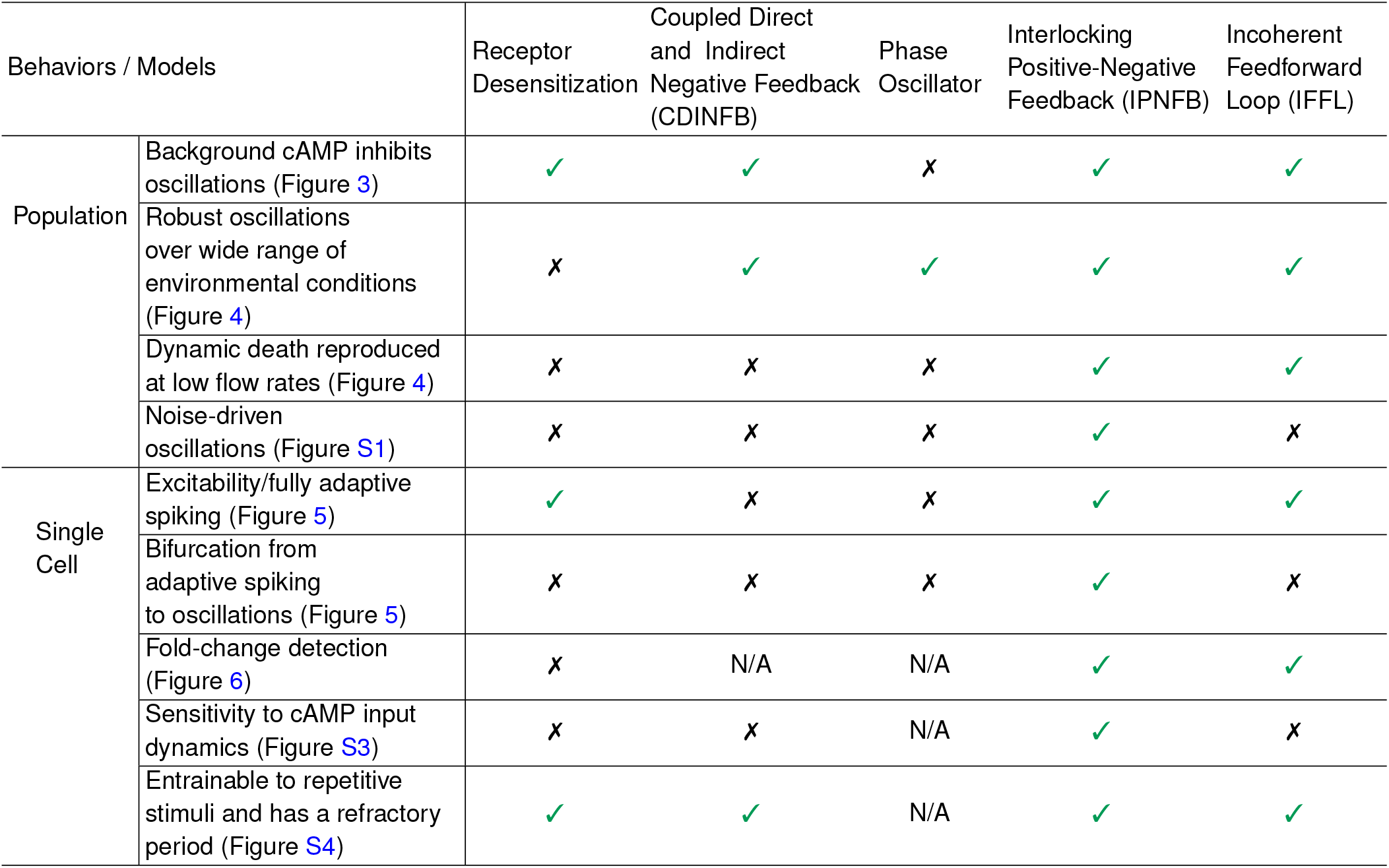
Population and single-cell behaviors evaluated in different models. Green check marks indicate if the model successfully reproduces the specific experimental behavior. X marks indicate the model is unable to reproduce the experimental behavior. N/A (not applicable) indicates the experimental comparison cannot be made because the model either fails to produce single-cell adaptive spikes (as for the Phase Oscillator model) or only shows partial adaptation (as for the CDINFB model).

| Behaviors / Models |  | Receptor Desensitization | Coupled Direct and Indirect Negative Feedback (CDINFB) | Phase Oscillator | Interlocking Positive-Negative Feedback (IPNFB) | Incoherent Feedforward Loop (IFFL) |
| --- | --- | --- | --- | --- | --- | --- |
| Population | Background cAMP inhibits oscillations (Figure 3) | ✓ | ✓ | ✗ | ✓ | ✓ |
|  | Robust oscillations over wide range of environmental conditions (Figure 4) | ✗ | ✓ | ✓ | ✓ | ✓ |
|  | Dynamic death reproduced at low flow rates (Figure 4) | ✗ | ✗ | ✗ | ✓ | ✓ |
|  | Noise-driven oscillations (Figure S1) | ✗ | ✗ | ✗ | ✓ | ✗ |
| Single Cell | Excitability/fully adaptive spiking (Figure 5) | ✓ | ✗ | ✗ | ✓ | ✓ |
|  | Bifurcation from adaptive spiking to oscillations (Figure 5) | ✗ | ✗ | ✗ | ✓ | ✗ |
|  | Fold-change detection (Figure 6) | ✗ | N/A | N/A | ✓ | ✓ |
|  | Sensitivity to cAMP input dynamics (Figure S3) | ✗ | ✗ | N/A | ✓ | ✗ |
|  | Entrainable to repetitive stimuli and has a refractory period (Figure S4) | ✓ | ✓ | N/A | ✓ | ✓ |

#### 1. Population Behaviors

##### I. Sustained population cAMP oscillations and their repression by background cAMP application

The most distinctive signaling behavior *Dictyostelium* displays is synchronized cAMP oscillations across populations. These oscillations are mediated by cell-cell communication through a relay mechanism where cells produce cAMP internally upon detecting elevated levels of external cAMP. All of the models investigated recapitulate this phenomenon of synchronized population oscillations (Figure 3A, before external cAMP input). More recently, experiments demonstrate these collective oscillations can be suppressed by suddenly increasing the concentration of external cAMP (Figure 3A) [8]. All the models except for the Phase Oscillator model reproduce this behavior – the Phase Oscillator model displays the opposite behavior with higher background cAMP levels leading to faster and more coherent group oscillations (Figure 3B-F). This discrepancy suggests the observed population oscillations do not arise from a mechanism where coupled individual cells oscillate autonomously even in the absence of external cAMP.

**FIG. 3:**
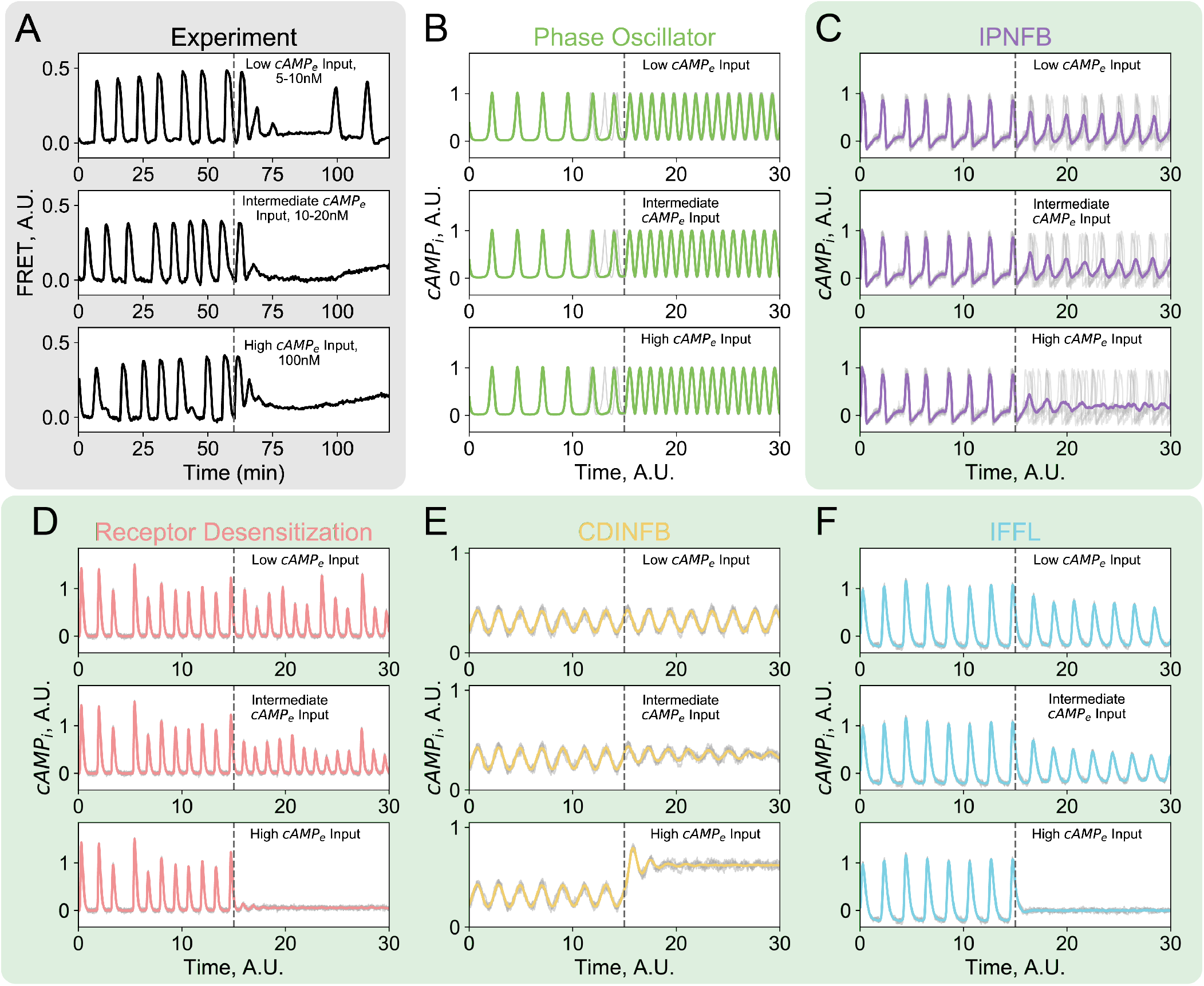
Evaluating models against population phenomena: population oscillations are suppressed by a step input of external cAMP. **(A)** Experimental data adapted from Sgro, et al. [8] shows population-wide oscillations slow down upon low and intermediate external cAMP addition, and are completely suppressed by high concentrations of external cAMP. **(B)** The Phase Oscillator model does not reproduce the experimental data. The rest of the models investigated all qualitatively reproduce the experimental data, despite acting through two distinctive mechanisms. **(C)** In the IPNFB model, population oscillations slow down due to desynchronization between single cells upon external cAMP addition. **(D F)** In the other three models, population oscillations slow down through inhibition of adaptive spike height in single cells. For model simulation results, solid colored traces are the mean internal cAMP of all the cells in the population, and gray traces represent single-cell dynamics from five cells in the population. The low, intermediate, and high cAMP levels in each model are determined arbitrarily, with a low external cAMP level slightly suppressing oscillations and a high level inhibiting the population oscillations entirely, except in the Phase Oscillator model as its oscillations speed up with increasing external cAMP. Gray dashed lines indicate the start of the step input of external cAMP. A gray shaded background highlights experimental data. A green shaded background indicates models that reproduce the experimental observations.

In the remaining models, group oscillations are suppressed by background cAMP because the added cAMP masks the cell-secreted external cAMP that propagates through the population. However, this inhibition of population oscillations occurs through two distinct mechanisms depending on the model. One mechanism, exploited in the IPNFB model, is that raising background cAMP tunes the single-cell spiking rate and decreases the coherence of single-cell spiking events. As a result, the amplitude of the single-cell spikes remains constant (Figure 3C, see gray traces for single cells) and the loss of collective oscillations is due to desynchronization between cells. The other mechanism, common to the remaining models, is that increasing background cAMP concentrations still result in coherent population-level oscillations, but now with a reduced oscillation amplitude. This can be seen in the simulation data where at intermediate background cAMP levels, single cells within the population still oscillate coherently, while the spike heights are reduced (Figure 3D - F, see gray traces for single cells). It should be possible to distinguish between these two mechanisms in future experiments by more extensively analyzing single-cell behaviors within populations in response to the addition of external cAMP.

##### II. Population oscillations depend on environmentally-mediated cell-cell coupling

Collective oscillations in *Dictyostelium* are mediated by the signaling molecule cAMP in a shared media. As a result, population-level oscillations depend strongly on environmental cell-cell coupling parameters such as cell density and the cAMP degradation rate (either through native enzymatic means using phosphodiesterases or through physical means such as fluid flow around the cells). How these parameters coordinate the emergence of population-wide oscillations can be experimentally explored by varying cell density and altering media flow rates over cells. Previous experimental work demonstrates that in most regimes, group oscillations emerge as cell-cell coupling strength increases through increasing cell seeding density or decreasing media flow rate. (Figure 4A) [7].

**FIG. 4:**
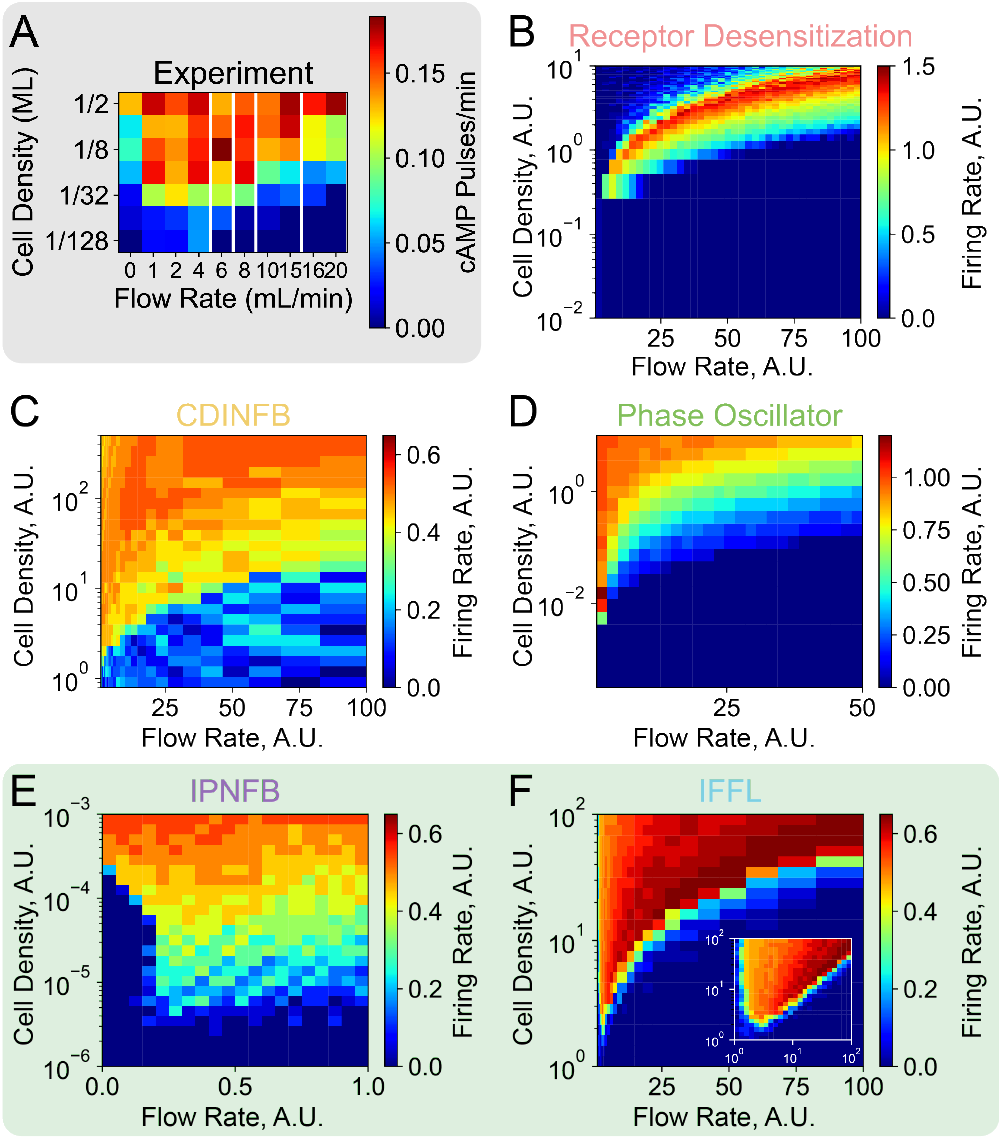
Evaluating models against population phenomena: population oscillations depend on cell density and external cAMP media flow rate. **(A)** Experimental population firing rate phase diagram for *Dictyostelium* cells in a perfusion chamber with varying media flow rates and cell densities measured in fractions of a monolayer (ML) from Gregor, et al. [7]. **(B)** Unlike the experimental data, the Receptor Desensitization model only displays population oscillations in a narrow parameter range. **(C and D)** The CDINFB and Phase Oscillator models successfully reproduce collective oscillations over a large range of parameter space, but do not capture the dynamic death region at low flow rates found in the experimental data. **(E and F)** The IPNFB and IFFL models perform the best at reproducing the experimental data because both models display robust oscillations over a large parameter space as well as display a dynamic death regime at low flow rates. Inset in **(F)** shows simulation results plotted with a logarithmic x-axis to highlight the dynamic death regime in the IFFL model. We added noise to single cells in the population simulations as detailed in the *Materials and Methods*. A gray shaded background highlights experimental data. A green shaded background indicates models that reproduce the experimental observations.

All of the models investigated qualitatively recapture the emergence of group oscillations as cell-cell coupling increases (Figure 4B-F). However, there are two key experimental features of the observed coupled oscillations that can be used to distinguish between models. First, these population-wide oscillations are experimentally observed across a large coupling parameter regime (Figure 4A). This observation is not reproduced by the Receptor Desensitization model, which displays collective oscillations only in a narrow regime (Figure 4B). The underlying reason is that sustained oscillations in this model are possible only when external cAMP concentrations are restricted to a narrow dynamic range where the receptors are not saturated. Second, in the region where the external flow rate is extremely low, the population oscillations die off at all but the highest cell densities, due to a phenomenon known as ”dynamic death” (Figure 4A, 1 mL/min Flow Rate). Past theoretical modeling efforts suggest dynamic death exists because external medium dynamics are too slow to catch up with the faster internal cAMP dynamics [19, 20]. In model simulations, only the IPNFB and IFFL models recapture this experimental observation (Figure 4E and inset in F).

Two major studies of population oscillations in *Dictyostelium* suggest a major role for noise in initiating and maintaining population oscillations [7, 8]. For this reason, we assessed how including noise alters population-level oscillations in each model by constructing stochastic generalizations of models without noise (Figure S1). Simulation results show that for the Receptor Desensitization, CDINFB, and IFFL models, including noise in the single cell networks leads to a more gradual transition between no population-wide oscillations and collective oscillations but no other qualitative changes in behavior (Figure S1B, C, F). Both the Phase Oscillator and IPNFB models, which were designed with stochasticity, display a change in behavior without noise. In the Phase Oscillator model, both in the original study and in this new analysis, noise aids in the initiation of population oscillations by lowering the critical (lowest) cell density that allows for the oscillations (Figure S1D) [7]. However, in the IPNFB model, noise is required for initiating and coordinating population oscillations because the removal of noise completely disables population-wide oscillations (Figure S1E), suggesting noise is critical for driving population behaviors in the IPNFB model [8].

#### 2. Single-Cell Behaviors

##### I. Single cells bifurcate from adaptive spiking to sustained oscillations in response to external cAMP step input

One of the most prominent experimentally-observed behaviors in isolated single *Dictyostelium* cells is that a small step change in external cAMP concentrations results in a large, sudden increase of internal cAMP concentration, which then returns back to near-baseline levels [7, 8]. This response has been termed an adaptive spike as it is reminiscent of similar phenomena in neural systems (Figure 5A). Experiments further show that if even higher levels of external cAMP are applied, single cells abruptly switch their response from producing an adaptive spike to an adaptive spike followed by sustained oscillatory behavior where internal cAMP concentrations oscillate in time even though external cAMP concentrations are constant (Figure 5B) [8]. These experiments suggest that external cAMP concentrations serve as a “bifurcation parameter” that drives the *Dictyostelium* signaling networks across a bifurcation from an excitable to an oscillatory regime.

**FIG. 5:**
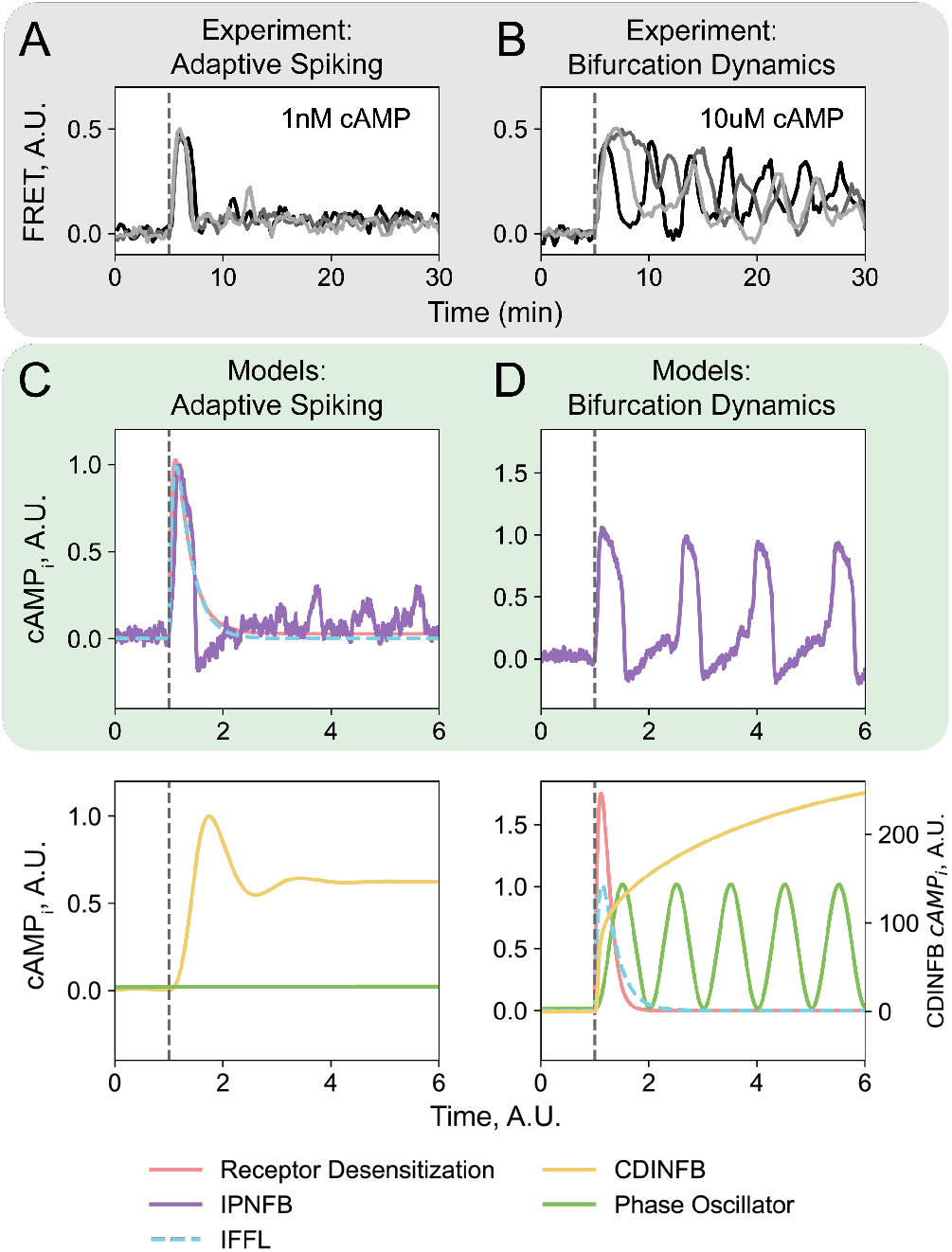
Evaluating models against single cell phenomena: cells bifurcate from adaptive spiking to sustained oscillations in response to external cAMP step input. **(A and B)** Experimental data shows single *Dictyostelium* cells display a bifurcation in response to an external cAMP concentration step input, from adaptive spiking **(A)** to sustained oscillations **(B)** (n = 3 example cells per condition). Data is adapted from Sgro, et al. [8]. **(C and D)** Model simulations with low and high external cAMP step inputs show that only the IPNFB model recapitulates both behaviors, with the green-shaded upper panels displaying models that recapitulate the experimental observations, specifically the adaptive spiking behavior **(C)** and the bifurcation to sustained oscillations **(D)** and the lower panels showing models that do not recapitulate these phenomena. In both the experimental data and model results, external cAMP is applied at the gray dashed line.

To identify which network architectures could reproduce the key experimental observations, we ran simulations to check if the models produced: (1) single, excitable spikes after a low-level external cAMP step input and (2) a spike followed by a bifurcation to oscillations after a high-level cAMP step input. The Receptor Desensitization model, the IPNFB model, and the IFFL model produce canonical adaptive spikes where internal cAMP levels return back to near-baseline levels after a large response (Figure 5C). In contrast, the CDINFB model produces a large spike in response to a small input but fails to return back to near-baseline levels, and the Phase Oscillator model fails to show any sort of adaptive spiking dynamics (Figure 5C). For this reason, for the remainder of the single cell phenomena tests beyond exploring the response to a large step input of external cAMP, we focus on the three models that produced adaptive spiking, as this is critical for all singlecell level phenomena. Further simulation results on the other models are in the *Supplementary Materials*.

Next, we simulated single cell responses to large step inputs of external cAMP, with a specific interest in whether or not the the three models that show adaptive spikes also show a bifurcation to oscillatory behavior at high levels of external cAMP. The height of this large step input was chosen to be 10^4^ times the low concentration based on experimental observations in Sgro, et al. that show robust oscillations at 10 μM external cAMP [8]. Only the IPNFB network displayed the bifurcation from excitable adaptive spiking to sustained oscillations (Figure 5D). The bifurcation to oscillations in the IPNFB model proceeds through a standard supercritical Hopf bifurcation based on a negative feedback loop with a time delay [21, 22]. Mechanistically, the origin of these oscillations is that the activator must build up to a sufficiently high level to activate the inhibitor, which then turns off activator production. The other two models lack either negative feedback (IFFL model) or a time delay (receptor desensitization only depends on instantaneous cAMP levels), accounting for the lack of single cell oscillations (Figure 5D). However, as discussed above, these later models can still support sustained population-level oscillations as the population-level oscillations originate from synchronized adaptive spikes in single cells.

##### II. Fold-change detection of external cAMP levels

During development, robust collective cAMP oscillations are observed from approximately 4 hours until as late as 20 hours post-starvation [7]. During this process, external cAMP levels vary dramatically because of changes in cell density due to cell migration. Single cells must robustly respond to this constantly changing and noisy external signal. Recent experiments show that single *Dictyostelium* cells conduct fold-change detection [17]: the response of single cells is determined only by the relative change in input signal (i.e. the ratio of the final cAMP concentration to the initial cAMP concentration), rather than absolute changes (Figure 6A). An important feature of fold-change detection is that it allows cells to operate in similar ways across a wide range of background cAMP concentrations [23]. This suggests that fold-change detection may underlie the ability of the *Dictyostelium* signaling networks to function in spatially and temporally heterogeneous environments during the development process.

**FIG. 6:**
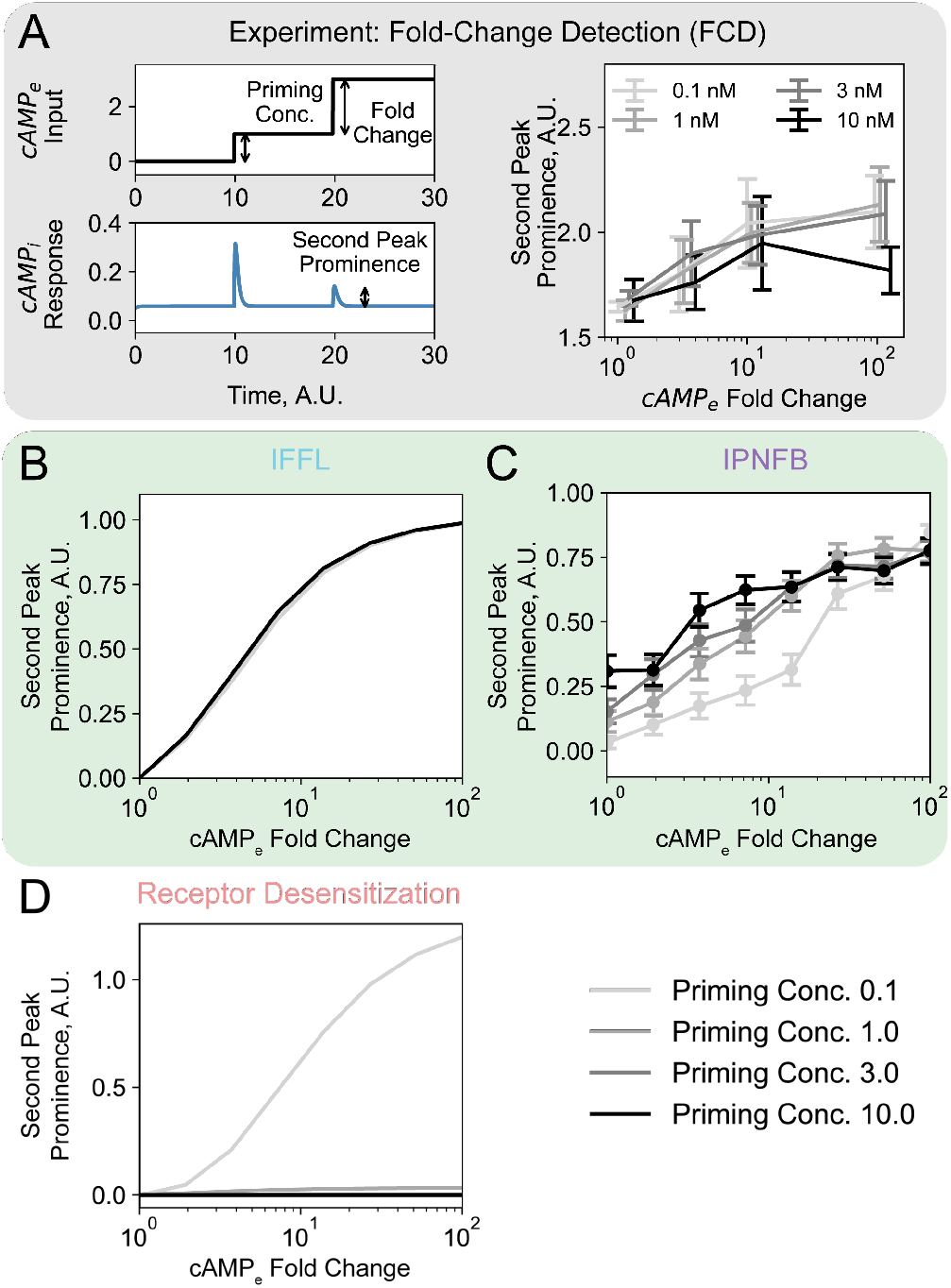
Evaluating models against single cell phenomena: cells are sensitive to fold changes in external cAMP levels. **(A)** Experimental data shows *Dictyostelium* cells conduct fold-change detection. Left panel: diagram depicting how fold-change detection is quantified in both experimental and simulation data. Cells experience two consecutive external cAMP step inputs, with the first step height being the “Priming Concentration” and the fold change of the second step height over the first step height being the “Fold Change”. The height of the internal cAMP spike in response to the second step input is quantified as “Second Peak Prominence” throughout the figure. Right panel: experimental data adapted from Kamino, et al. [17] shows single cells exhibit fold-change detection. **(B D)** Simulation of different model responses shows that only the IPNFB and IFFL models qualitatively reproduce fold-change detection **(B and C)**. The IFFL traces representing different priming concentrations collapse on a single line in **(B)**. To account for single-cell noise in the IPNFB model, we ran 50 simulations for each priming concentration-fold change pair and display the mean and standard error of the mean from the simulations in **(C)**. The Receptor Desensitization model response traces to 3 and 10 unit priming concentrations collapse on a single line **(D)**. In simulations the one unit priming concentration is set to match the “low” *cAMP*_*e*_input level in Figure 5. A gray shaded background highlights experimental data. A green shaded background indicates models that reproduce the experimental observations.

To measure fold-change detection capabilities in the three models that reproduce single-cell adaptive spikes, we ran simulations that mimic the experimental design of Kamino, et al. where cells were subject to two consecutive step changes in external cAMP at different concentrations (the priming concentration and the secondary concentration, see Figure 6A) [17]. As in the experimental work, fold-change detection was measured by the prominence of the second adaptive spike and we scanned model responses to initial priming concentrations of 0.1 external units through 10 external units of cAMP. For a network that performs perfect fold-change detection, we expect the response curves for different priming concentrations to collapse on a single line, a phenomenon observed in experiments (Figure 6A). The model with the best fold-change detection capabilities is the IFFL model which shows nearly perfect, deterministic fold-change detection over almost two orders of magnitude changes in external cAMP levels (Figure 6B). The IPNFB model, which is built with noise, also displays approximate fold-change detection (Figure 6C). In contrast, the Receptor Desensitization model does not exhibit fold-change detection (Figure 6D).

Interestingly, the IFFL and the IPNFB models achieve fold-change detection through two very different mechanisms. In the IFFL model, the response to fold-differences with different step heights is identical [17]. In contrast, the IPNFB model achieves only approximate fold-change detection through a combination of logarithmic-sensing and stochasticity-mediated modulation of the single cell firing rate (see Figure 6C) and Figure S2).

#### 3. Relating Network Architecture to Collective Behaviors

Our unified simulation framework allows us to identify the key components and network features needed to reproduce specific experimental observations (Figure 7). The analyzed models include disparate design features such as negative and positive feedback control, logarithmic environmental sensing, incoherent feedforward loops, and modeling single cells as phase oscillators. Nonetheless, there are some common themes that emerge from our analysis.

**FIG. 7:**
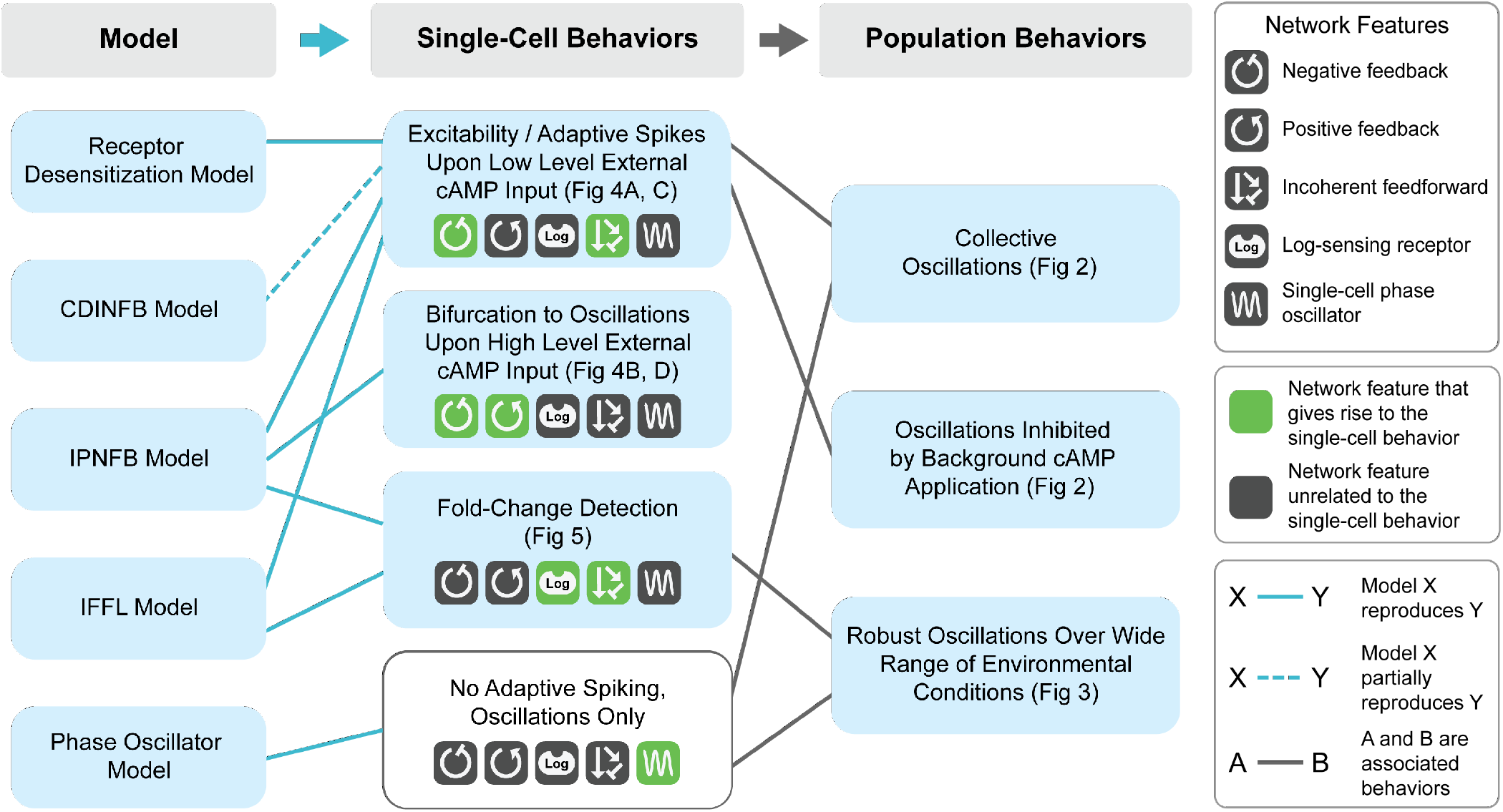
Graphical summary of our analysis reveals which network features give rise to single-cell and population-level experimental behaviors. All models except the Phase Oscillator model partially or fully reproduce at least one of the experimentally-observed single cell behaviors. These single-cell behaviors are driven by different network features in the models, suggesting there are multiple ways cellular signaling network architectures can generate these behaviors. Furthermore, several of the single-cell behaviors are associated with population-level behaviors, suggesting that the network features underlying the associated single-cell behaviors are critical for driving the associated population-level behaviors.

First, we find that negative feedback or feedforward motifs appear to be especially critical to produce excitability (i.e. adaptive spiking) at the single-cell level. These two features appear in all of the models except the Phase Oscillator model, the only model that fails to reproduce adaptive spiking on the single-cell level. This suggests that negative feedback or incoherent feedforward motifs drive single-cell excitability, consistent with past theoretical findings that negative feedback and incoherent feedforward loops are the core topologies that drive adaptive spikes [24]. Similarly, all of the models other than the Phase Oscillator model reproduce the observation that background cAMP inhibits collective oscillations, suggesting single-cell excitability may be the driving mechanism for this behavior.

Another key design feature that seems to be important for coordinating *Dictyostelium* behavior is positive feedback control. Positive feedback is a key element of the IPNFB model, the only model of the five we analyzed that can completely recapitulate all experimental observations at the single-cell level. The idea that positive feedback control might facilitate the single-cell bifurcation to oscillations is again consistent with past theoretical work demonstrating that interlocking positive-negative feedback loops give rise to more robust and reliable oscillatory behaviors [25].

We also found that both incoherent feedforward loops – present in the IFFL model – and logarithmic sensing – present in IPNFB model – enable fold-change detection [26, 27]. This observation, along with the fact that only these two models reproduce this phenomenon, suggests that one of these design features is necessary to drive fold-change detection in this context.

Finally, we find that in order to have robust population-level oscillations that persist despite environmental variations, single cells must either perform fold-change detection or behave as a phase oscillator. Given the inability of phase-oscillator models to recapitulate single-cell behavior, our analysis suggests that cells likely implement fold-change detection in order to coordinate population behaviors.

## II. DISCUSSION

Many different properties of single cells and cellular populations have been proposed to be conceptually important for coordinating collective oscillations in multicellular systems, including *Dictyostelium*. Here, we exploited the stereotypical spiking behavior displayed by single *Dictyostelium* cells to normalize models and directly compare them to each other, as well as to experimental observations. Our results show two types of networks fully describe the population-level behaviors: the IPNFB with a logarithmic pre-processing module and the Incoherent Feedforward Loop (IFFL) (Table I). These networks both display two key single-cell level design features that are critical for coordinating the observed population-wide dynamics: excitable single-cell level dynamics and foldchange detection (Table I, Figure 7). Furthermore, only the IPNFB model reproduces all of the observed singlecell and population-level behaviors. This suggests that while both networks are potential pathways for the design of collective oscillatory systems, the IPNFB model more accurately reflects the specific *Dictyostelium* signaling network. Through our analysis, we find that the observed population-level phenomena can be driven by a key single cell network design principle: excitable dynamics that respond to relative changes in external signals.

Excitability is a critical single-cell network design feature that drives collective oscillations that are tunable by cellcell coupling (Figure 7). Molecular networks that display excitable dynamics require two network features. The first feature is that the network includes either a negative feedback or incoherent feedforward motif that brings down the levels of or activities of the molecular species that are activated by external inputs [24]. Negative feedback and incoherent feedforward loops that drive singlecell excitability are found in the two models that succeed in recapitulating almost all experimental observations (Table I). The second feature aiding excitable dynamics is timescale separation/time delay where after the increase of the activator species, the inhibitory interactions either respond on a slower timescale or have a built-in delay. This allows the input activation to dominate the early response before a strong inhibitory response begins, leading to an excitable spike in the activator species. In both example networks with superior performance, there is a time separation between the inhibitor species *τ*_*I*_ and the activator species *τ*_*A*_ facilitating excitability in the networks (see Figure S5 for the effect of decreasing timescale separation). This timescale separation is naturally accompanied by refractoriness as is observed experimentally in single *Dictyostelium* cells (Table I, Figure S4). This refractoriness leads to unidirectional signal propagation in a population.

The second critical single-cell network design feature we identified is that internal dynamics must be dependent on the relative change of external signal as opposed to absolute concentrations, a feature known as foldchange detection (Figure 7). This feature coordinates robust population oscillations during development over a large range of environmental parameters such as large changes in cell density, varying ability to degrade external cAMP, and noisy fluctuations (Figure 4). Previous work has identified several major classes of fold-change detection-competent networks by conducting exhaustive scans of network topologies [26, 27]. These scans found fold-change detection-competent models are extremely rare, and the two naturally occurring fold-change detection models, the incoherent type I feedforward loop (I1FFL) and the non-linear integral feedback loop (NLIFL), are among the simplest networks suggested to achieve optimal response amplitude, speed, and noise-resistance. Another possible mechanism for achieving fold-change detection is logarithmic sensing at the receptor level which can theoretically be achieved with an allosteric protein [28]. The best-performing example networks from our analysis both fall into the canonical categories: one is an IFFL network and the other uses a logarithmic-sensing module (Figure 1 and 6). Although both networks display fold-change detection, we note there are subtle differences. With the IFFL model, the network tunes spike height in response to input fold change. The IPNFB model, however, modulates the probability of spikes in individual cells in response to input fold-change (Figure S2), tuning the average response of a group of cells and thus displaying fold-change detection at the population level.

Additionally, our work also further reinforces previous findings that stochastic noise in the single cell network potentially plays a role in coordinating population behaviors. Previous studies on bacterial competence suggests that noise combined with an excitable module can explain both the initiation of and escape from the competent state, and that noise levels modulate the percentage of cells entering into and exiting from the competent state in bacterial communities [29, 30]. In *Dictyostelium*, work on the cAMP signaling and chemotaxis networks suggests that noise in network components with both positive and negative feedback loops plays a vital role in coordinating network dynamics [8, 31]. In our analysis, in the model that most accurately recapitulates behaviors in *Dictyostelium*, noise is crucial for fold-change detection and coordinating population oscillations (Table I, Figure S1). Stochasticity could provide a mechanism for initiating and maintaining population-wide oscillations without requiring specialized cells such as pacemaker cells for robustly coordinating the population, allowing for decentralized control. To answer whether and how noise modulates population-wide behaviors, better experimental techniques and longer timescale experiments that quantify noise-driven phenomena and modulate potential sources of noise are required. Taken together, our analysis suggests that excitability that is insensitive to absolute environmental signals acts as a key network design feature that allows for efficient oscillations coordination in *Dictyostelium* populations, and noise in signaling networks plays a role in population-behavior modulation.

Coordinated population behaviors not only require specific features in single-cell signaling networks, but also require coupling between cells. When the cells are coupled through the external medium, the timescales of the internal and external signal dynamics need to be effectively coupled. In the case of *Dictyostelium* populations, the time scale of the external signal dynamics is modulated by cAMP degradation and this rate is determined by phosphodiesterase concentrations and kinetics. At extremely high cAMP degradation rates, single cells cannot effectively communicate. By contrast, if the degradation rates are too low, the dynamics of the medium are too slow to follow the internal dynamics, leading to desynchronization of single cells, or “dynamic death” [19, 20], a phenomenon experimentally observed in Gregor, et al. and Sgro, et al. [7, 8]. In our analysis, the IPNFB and IFFL models reproduce the dynamics death phase, suggesting that cells match the timescales governing internal signaling networks and external signal propagation. Such timescale coordination may also be crucial for chemotaxis in *Dictyostelium* cells [32].

While all of the models investigated here take a mean field approach – neglecting space and time for signal propagation between cells – in reality signals propagate though the population and create complex spatiotemporal patterns such as concentric waves and rotating spirals [6]. Traveling waves are a natural analogue of the kind of coherent population level oscillations discussed in this work and may represent an important design principle for coordinating behavior across large spatial reasons. In support of this idea, we note that waves are also observed during development, where mouse embryonic cells use excitable internal YAP dynamics that are coupled by Notch signaling to achieve long-range oscillatory waves for vertebrate segmentation [33], and *Drosophila* embryos use propagating Cdk1 waves to synchronize cell cycles across large spatial scales [34]. In synthetic biology, engineered negative and positive feedback motifs can achieve excitability and robust waves across a large bacterial population when coupled through diffusible molecules [35, 36].

Altogether, this work identifies excitability and foldchange detection as key design features of internal signaling networks that allow for robust coordination of population-wide oscillations in heterogeneous ill-defined environments. These network features found in *Dictyostelium* are widely shared in natural and synthetic cell populations that display collective oscillatory behaviors, suggesting these network features could be a common control mechanism used by biological systems to coordinate signal transduction in multicellular contexts. These features enable many desirable behaviors for multicellular populations including coordination over orders of magnitude differences in population size, environmental insensitivity to small fluctuations, and fast, unidirectional long-range signal propagation. Our work suggests that there are relatively few signaling network design motifs that can robustly coordinate emergent multi-scale behaviors in biological systems.

## III. MATERIALS AND METHODS

Model equations were adapted from the literature [7, 8, 14, 16, 17], and the mathematical expressions and parameters for each model are detailed in the *Supplementary Materials*. All simulations were solved by the Euler-Maruyama method. Time step sizes were empirically chosen to make sure simulation outputs are reliable such that decreased time step sizes would not produce alternative results. Exploring the emergence of population oscillations and their dependence on cell-cell coupling (Figure 4) requires investigating a wide range of parameter space and thus smaller time step sizes were taken to make sure the simulation outputs are reproducible. Models were normalized to one another as described in the *Results* section.

Noise in single-cell networks was added as a Langevin noise term *η*(*t*) to the end of the equation representing the internal “activator” component. The noise term satisfies *< η*(*t*)*η*(*t*′) *>*= *σ*^2^*δ*(*t* − *t*′), with *σ*^2^ denotes the noise strength. The noise strength in each model was chosen either from the original literature for models with noise in the original implementations, specifically the Phase Oscillator model and IPNFB model, or arbitrarily determined such that noise allows for a slight quantitative change in the phase diagram describing the emergence of population oscillations but not a qualitative change (Figure S1). The respective noise strengths in the Receptor Desensitization model, CDINFB model, Phenomenological Phase Oscillator model, IPNFB model, and IFFL model were set to 10, 0.1, 0.02, 0.15, 0.01.

## Code Availability

Experimental data and Python code for all simulations are available on GitHub at https://github.com/sgrolab/dictymodels.

## ACKNOWLEDGMENTS

The authors thank Dr. Hernan Garcia, Dr. Emily Hager, Dr. Chiara Ricci-Tam, Mark Aronson, Noshin Nawar, and Breanna O’Reilly for their critical reviews of our manuscript and helpful comments. This work was supported by the National Institutes of Health (NIGMS) grant R35 GM133616 to A.E.S., the Burroughs Wellcome Fund Career Award at the Scientific Interface to A.E.S., the National Institutes of Health (NIGMS) grant R35 GM119461 to P.M., and a Simons Foundation MMLS grant to P.M. X.Z. was partially supported by the Fonds de recherche du Québec Nature et technologies (FRQNT).

