## Supplementary Materials for "Robust coordination of collective oscillatory signaling requires single-cell excitability and fold-change detection"

### A. Model Details and Assumptions

#### I. Receptor Desensitization model

One of the earliest and most widely used models of population-level cAMP oscillations in *Dictyostelium* was developed by Martiel and Goldbeter [14] in response to the experimental observation that the cAMP receptor CAR1 is desensitized as external cAMP concentration increase [37, 38]. In this model, cells detect external cAMP using the cAMP receptor CAR1 ($R$ in Figure 1), leading to the production and release of internal cAMP into the environment. As external cAMP concentrations increase, a negative feedback loop is triggered that leads to receptor desensitization and a corresponding decrease in internal cAMP production and excretion. This well-understood mechanism based on a negative feedback loop (i.e. receptor desensitization) with a time delay that naturally leads to population-level oscillations [39].

#### II. Coupled Direct and Indirect Negative Feedback (CDINFB) model

The authors of Maeda, et al. [16], building on earlier works including the Laub-Loomis model [15], developed a detailed mechanistic model of the *Dictyostelium* signal relay network. In this model, external cAMP binds to the CAR1 receptor ($R$ in Figure 1), leading to production of internal cAMP by the enzyme adenylyl cyclase (ACA, shown as $A$ in Figure 1). The activation of ACA also triggers a pair of coupled negative feedback loops: a direct negative feedback mediated through protein kinase A (PKA, shown as $I$ in Figure 1) that turns off cAMP production, and an additional indirect negative feedback mediated by the proteins RegA and ERK2 that activates PKA, once again leading to decrease in cAMP production. Additionally, the ERK2-RegA path leads to internal cAMP decrease through RegA acting as a internal phosphodiesterase that degrades cAMP.

#### III. Phenomenological Phase Oscillator model

Inspired by new experiments showing that even isolated single cells can oscillate in response to elevated levels of external cAMP, the authors of Gregor et al. [7] constructed a phenomenological model inspired by the Kuramoto model [40, 41] where the *Dictyostelium* signaling network is modeled as a phase-oscillator, with the phases of different cells coupled through the external cAMP concentration. Internal cAMP concentrations are assumed to be directly proportional to the $sine$ of the phase of the oscillator.

#### IV. Interlocking Positive-Negative Feedback (IPNFB) model

This phenomenological model introduced in Sgro, et al. and expanded on in Noorbakhsh, et al. [8, 20] captures the key behaviors of the *Dictyostelium* signaling network using a generalization of the FitzHugh Nagumo (FHN) model, which commonly used in neuroscience to model excitable neural dynamics [21, 22]. The model consists of two elements: an activator species ($A$) and an inhibitor species ($I$). The presence of external cAMP leads to increased production of the activator species, suggested to be representative of adenylyl cyclase activity, which can trigger a fast positive feedback loop that results in even more activator production. The activator also triggers a slower negative feedback by producing an inhibitor species that suppresses production of the activator, though this inhibitor species is suggested to be representative of a larger network structure and not a specific molecular species. In this model, internal cAMP concentrations are assumed to track the activator concentration, with cells releasing cAMP into the environment when the activator is above a certain threshold.

#### V. Incoherent Feedforward Loop (IFFL) model

Like the IPNFB model, the incoherent feedforward loop (IFFL) model is a phenomenological model of the *Dictyostelium* signaling network that combines fast activation with inhibition on longer time scales inspired by the experimental observations in Kamino, et al. [17]. In the IFFL model, the receptor up-regulates both the activator ($A$) as well as the inhibitor species ($I$). Crucially, while the inhibition varies linearly with the concentration of external cAMP, the activator exhibits a Hill-like (sigmoid-like) kinetics in response to external cAMP concentrations. This model displays interesting behaviors in the single-cell context including fold-change detection [23, 27] and perfect adaptation [24].

VI. Model assumptions
Finally, we note that an important but subtle difficulty involved in comparing and contrasting these models with experiments is that they make very different assumptions about how the models are related to experimental observables. Model parameters are often specified in arbitrary units, so it is *a priori* unclear how to relate predictions across the five models. To address this problem, we chose to non-dimensionalize both the amplitudes and timescales of internal cAMP responses in terms of the height and width of the adaptive spike induced by one unit of external cAMP input in single cells. In almost all of the models cells respond to a small, sudden change in external cAMP by producing a spike of internal cAMP. The height of this spike in each model gives a natural scale for measuring internal cAMP amplitudes whereas the width represents a natural time scale for normalizing times between models (see *Materials and Methods* for details).

#### B. Model Equations and Parameters

Note: Variable names in this section are as presented in the original literature. Some variable names in Figure 1 are renamed in order to emphasize which species act as different network components that may be common between models.

I. Receptor Desensitization model

The Receptor Desensitization model [14] is composed of three differential equations. $\rho_{T,i}$ and $\beta_i$ denote the proportion of receptors in the active state and internal cAMP concentration in the $i$th cell,and $\gamma$ stands for external cAMP concentration.

$$\frac{d\rho_{T,i}}{dt} = -f_1(\gamma)\rho_{T,i} + f_2(\gamma)(1-\rho_{T,i}) \tag{1}$$

$$\frac{d\beta_i}{dt} = q\sigma\Phi(\rho_{T,i},\gamma,\alpha) - (k_i + k_t)\beta_i \tag{2}$$

$$\frac{d\gamma}{dt} = k_t/h\frac{\sum^N \beta_i}{N} - k_c(\gamma - [cAMP]_{e,in}) \tag{3}$$

with:

$$f_1(\gamma) = \frac{k_1 + k_2\gamma}{1+\gamma}, \tag{4}$$

$$f_2(\gamma) = \frac{(k_1L_1 + k_2L_2c\gamma)}{1+c\gamma} \tag{5}$$

$$\Phi(\rho_{T,i},\gamma,\alpha) = \frac{\alpha(\lambda\theta + \varepsilon Y^2)}{1+\alpha\theta+\varepsilon Y^2(1+\alpha)} \tag{6}$$

$$Y = \frac{\rho_{T,i}\gamma}{1+\gamma} \tag{7}$$

$[cAMP]_{e,in}$ stands for cAMP concentration that is added externally and this notation is used across all models.

Parameter values are from Table II of the original literature[14]. Specifically, $k_1 = 0.036$, $k_2 = 0.666$, $L_1 = 10$, $L_2 = 0.005$, $c = 10$, $\lambda = 0.01$, $\theta = 0.01$, $\varepsilon = 1$, $q = 4000$, $k_i = 1.7$, $k_t = 0.9$,$k_c = 5.4$, $h = 5$, $N = 100$.

In Figure 4B, cell density and external cAMP degradation rates are varied by altering $\frac{1}{h}$ and $k_c$ .

Time normalization parameters for the Receptor Desensitization model is 6.94, and its height normalization parameter is 210.53.

II. Coupled Direct and Indirect Negative Feedback (CDINFB) model

The CDINFB model represents a signaling network consisting of seven molecular species [16]. Within a population of $N$ cells, respective concentrations of the biochemical species in the $i$th cell follow:

$$\frac{d[ACA]_i}{dt} = k_1[CAR1]_i - k_2[ACA]_i[PKA]_i \tag{8}$$

$$\frac{d[PKA]_i}{dt} = k_3[cAMP]_{cyt,i} - k_4[PKA]_i \tag{9}$$

$$\frac{d[ERK2]_i}{dt} = k_5[CAR1]_i - k_6[PKA]_i[ERK2]_i \tag{10}$$

$$\frac{d[RegA]_i}{dt} = k_7 - k_8[ERK2]_i[RegA]_i \tag{11}$$

$$\frac{d[cAMP]_{cyt,i}}{dt} = k_9[ACA]_i - k_{10}[RegA]_i[cAMP]_{cyt,i} \tag{12}$$

$$\begin{aligned}\frac{d[cAMP]_e}{dt} =& \rho k_{11}\frac{\sum^N [ACA]_i}{N} \\ & - k_{12}([cAMP]_e - [cAMP]_{e,in})\end{aligned} \tag{13}$$

$$\frac{d[CAR1]_i}{dt} = k_{13}[cAMP]_e - k_{14}[CAR1]_i \tag{14}$$

Parameters are $k_1 = 2, k_2 = 0.9, k_3 = 2.5, k_4 = 1.5, k_5 = 0.6, k_6 = 0.8, k_7 = 1.0, k_8 = 1.3, k_9 = 0.3, k_{10} = 0.8, k_{11} = 0.7, k_{12} = 4.9, k_{13} = 23, k_{14} = 4.5, \rho = 1, N = 100.$

In Figure 4C, cell density and external cAMP degradation rates are varied by altering $\rho$ and $k_{12}$ .

Time normalization parameters for the CDINFB model is 3.57, and its height normalization parameter is 3.15.

III. Phenomenological Phase Oscillator model

The Phase Oscillator model [7] considers each cell as a phase oscillator coupled by cAMP in the external medium. $[cAMP]cyt, i$ and $\theta_i$ denote the internal cAMP concentration and internal phase of the $i$th cell. Equations describing $[cAMP]cyt, i$ dynamics, $\theta_i$, and external cAMP $[cAMP]_e$ are:

$$[cAMP]_{cyt,i} = \frac{(-A_{max} + A_{bas})\sin\theta_i + A_{max} + A_{bas}}{2} \tag{15}$$

$$\frac{d\theta_i}{dt} = \omega(1 - \Phi([cAMP]_e)c_{excite} sin\theta_i) \tag{16}$$

With $\Phi([cAMP]_e) = \frac{K}{K+[cAMP]_e}$

$$\begin{aligned}\frac{d[cAMP]_e}{dt} =& \rho\frac{S_T V_C}{V_T S_C} c_{sec}\frac{1}{N}\sum^N [cAMP]_{cyt,i} \\ & - \frac{k}{Vt}([cAMP]_e - [cAMP]_{e,in})\end{aligned} \tag{17}$$

Parameter values are from the original paper [7]. Specifically, $A_{max} = 20, A_{bas} = 0.4, \omega = \pi/3, V_C = 1.1 \cdot 10^{-9}, S_T = 1.33, S_C = 1.3 \cdot 10^{-6}, K = 0.0004, c_{sec} = 3.6, c_{excite} = 1.01, N = 100, k = 5, \rho = 1/12, k = 5, V_t = 1.$

In Figure 4D, cell density and external cAMP degradation rates are varied by altering $\rho$ and $k$ .

Time normalization parameters for the Phase Oscillator model is 6, and its height normalization parameter is 19.6.

IV. Interlocking Positive-Negative Feedback (IPNFB) model

In the IPNFB model [8, 20], the signaling network in each cell is modeled following the FitzHugh-Nagumo framework [22]. The internal cAMP concentration and internal inhibitor concentration are represented by $A_i, R_i$. Equations describing $A_i, R_i$ and external $cAMP_e$ dynamics are:

$$\frac{dA_i}{dt} = A_i - \frac{1}{3}A_i^3 - R_i + I([cAMP]_e) \tag{18}$$

$$\frac{dR_i}{dt} = \varepsilon(A_i - \gamma R_i + c_0) \tag{19}$$

$$\begin{aligned}\frac{d[cAMP]_e}{dt} =& [cAMP]_{e,in} + \rho\alpha_0 + \rho S\frac{1}{N}\sum^N \Theta(A_i) \\ & - (J + \alpha_{PDE})[cAMP]_e\end{aligned} \tag{20}$$

With $I([cAMP]_e) = a \cdot log(1 + \frac{[cAMP]_e}{K_d})$, and $\Theta(A_i)$ being the Heaviside function: equal to 1 if $A_i >$ 0 and otherwise equal to 0. Parameter values are from the original paper [8]. Specifically, $\varepsilon = 0.1, \gamma = 0.5, c_0 = 1.2, N = 100, \alpha =$

$0.058, \alpha_0 = 800, \alpha_{PDE} = 1000, K_d = 10^{-5}, S = 10^6, \rho = 10^{-3.5}, J = 0.5$. An offset of 1.5 is applied to $A_i$ traces before outputs are height-normalized.

In Figure 4E, cell density and external cAMP degradation rates are varied by altering $\rho$ and $J$ .

Time normalization parameters for the IPNFB model is 27, and its height normalization parameter is 3.5.

V. Incoherent Feedforward Loop(IFFL) model

For the IFFL model [17], there are two key molecular players in each cell: an activator $y_i$ that approximates internal cAMP concentration and an repressor $x_i$ that represents internal inhibitor concentration. Equations describing $y_i, x_i$ and external $cAMP_e$ dynamics are:

$$\frac{dx_i}{dt} = \frac{1}{\tau}([cAMP]_e + \delta - x_i) \tag{21}$$

$$\frac{dy_i}{dt} = \frac{([cAMP]_e + \delta)^n}{([cAMP]_e + \delta)^n + (Kx_i)^n} - y_i \tag{22}$$

$$\frac{d[cAMP]_e}{dt} = \rho k_t y_i - \gamma([cAMP]_e - [cAMP]_{e,in}) \tag{23}$$

$\delta$ is included for lower detection limit. Parameter values are from the original paper [17]. Specifically, $\tau = 1.5, n = 2, K = 4, k_t = 2, \delta = 0.01, \gamma = 3, \rho = 0.01, N = 100$. An offset of 0.058 is applied to $y_i$ traces before outputs are height-normalized.

In Figure 4F, cell density and $cAMP_e$ degradation rates are varied by altering $\rho$ and $\gamma$.

Time normalization parameters for the IFFL model is 5.23, and its height normalization parameter is 0.26.

### SUPPLEMENTARY FIGURES

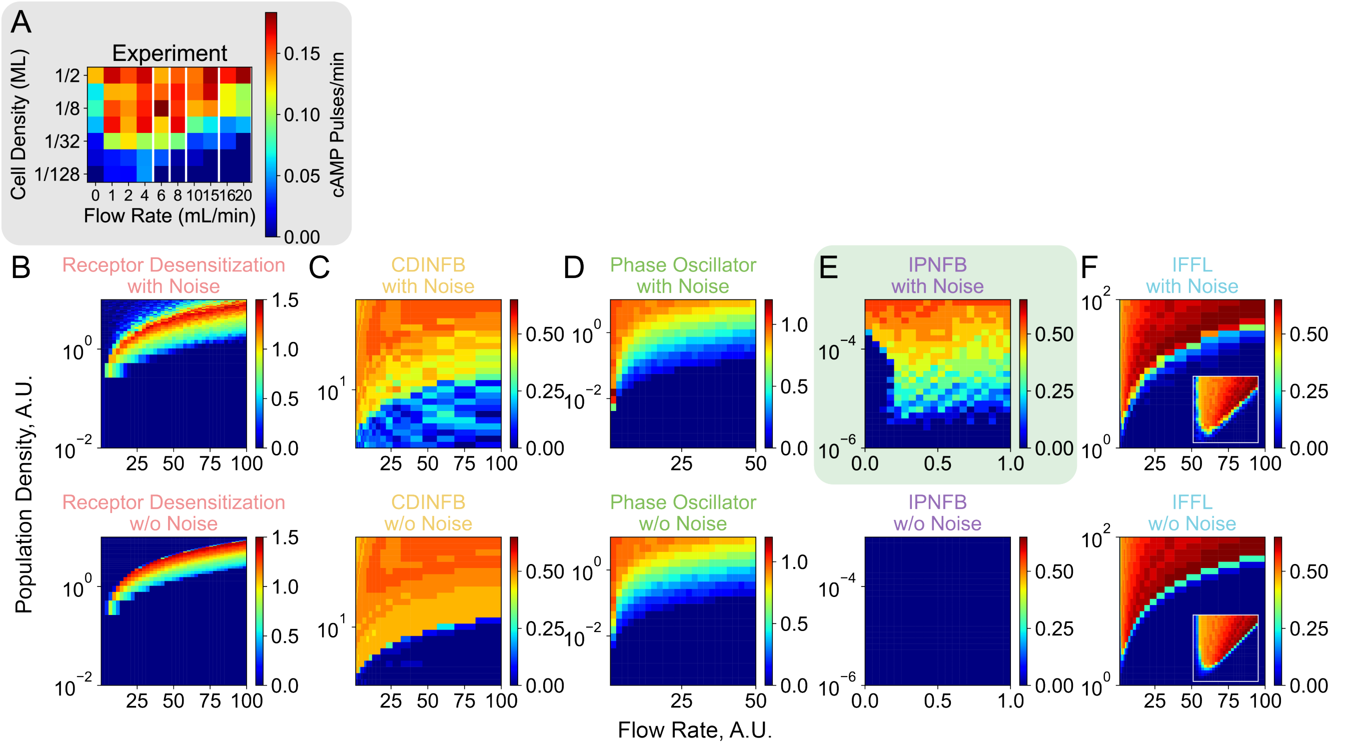


**FIG. S1: The role of noise in the emergence of population-wide oscillations. (A)** Experimental population firing rate phase diagram for *Dictyostelium* cells in a flow chamber with varying media flow rates and cell densities measured in fractions of a monolayer (ML) from Gregor, et al. [7]. **(B to F)**. Simulation results for all of the different models, with the upper panels displaying the results including single cell noise as shown in Figure 4, and the lower panels omitting single cell noise. Inset of **(F)** shows the same model output plotted with a logarithmic x-axis.

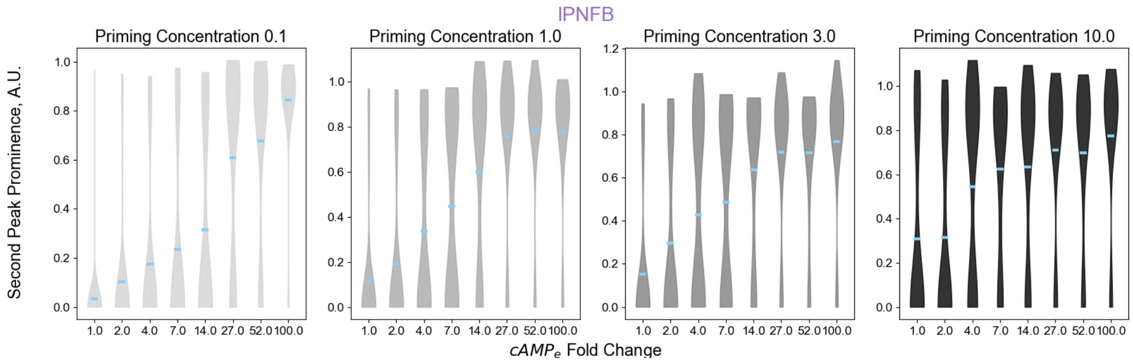


**FIG. S2: Probability density of IPNFB fold-change detection responses.** The IPNFB model has built-in single-cell noise so each simulation results in a different single-cell internal cAMP response. For each priming concentration-fold change pair, the means of the second-peak prominence (also shown in Figure 6C) are plotted as blue bars and the probability distributions are in shades of gray.

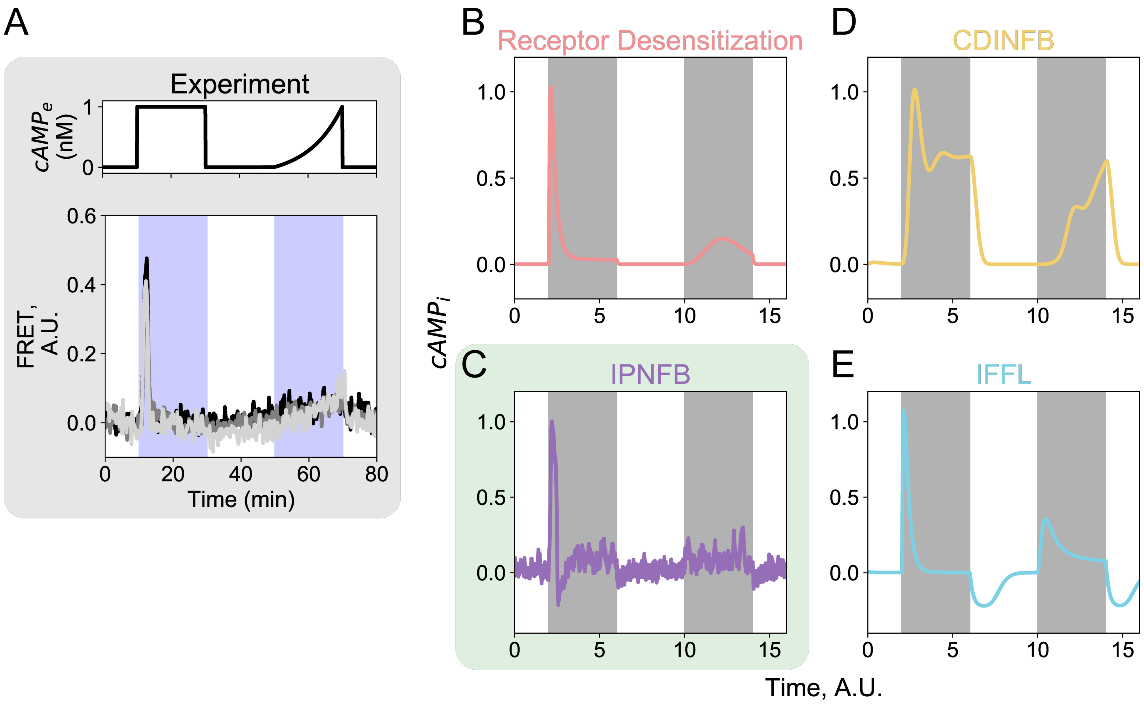


**FIG. S3: Different models have different internal cAMP responses to $[cAMP]_e$ input dynamics (A)** Experimental data show single *Dictyostelium* cells display an internal cAMP spike in response to an abrupt 1 nM step input of external cAMP but remain mostly quiescent in response to a slow ramp input to 1 nM external cAMP. Data adapted from Sgro, et al. [8]. Top panel: external cAMP input temporal profile. Bottom panel: Experimental $[cAMP]_i$ traces from three example cells. **(B - E)** Model simulations of cell responses. The first and second shaded region denotes when the step and exponential ramp $[cAMP]_e$ inputs are applied respectively. The maximum concentration of external cAMP input in each model is the same as in Figure 5C. A gray shaded background behind the plots highlights experimental data. A green shaded background indicates models that reproduce the experimental observations.

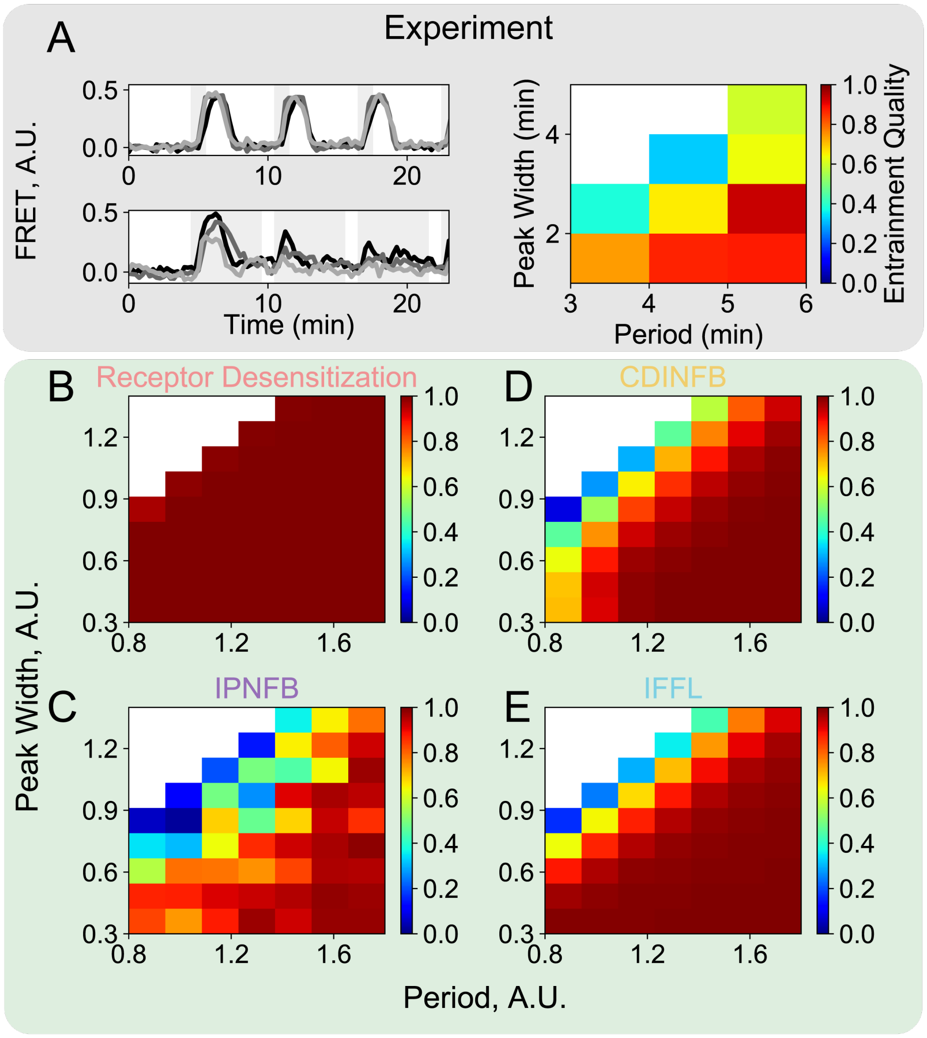


**FIG. S4: Entrainment quality varies with $[cAMP]_e$ input cycle period and input peak width. (A and B)** Experimental measurements adapted from Sgro, et al. [8] of how cells respond to repetitive external cAMP stimuli with different input peak widths and with different input cycle periods. This data illustrates single cells are entrainable and have a refractory period. Entrainment quality is quantified as the average Pearson's cross correlation coefficient of each of the subsequent $[cAMP]_i$ response spikes compared against the first response spike. **(A)** Left panel: $[cAMP]_i$ traces for three example cells experiencing a step input to 10 nM external cAMP. Shaded region indicates external cAMP input. Right panel: Entrainment quality increases with longer stimulation period and briefer peak width. **(B - E)** Model simulations show that all models that display adaptive spiking, even if not fully adaptive (Figure 5C), qualitatively match the experimental data. However, the entrainment quality metric does not take into account height variation between the first and subsequent spikes. The Receptor Desensitization model scores highly with this entrainment quality metric across the parameter regime we probe, but the spike height is reduced in some regimes indicating it has a refractory period (**B**). Based on what is used in experimental data, stimulation period and peak width is sampled from 0.8-1.8 and 0.4-1.4 times the respective timescale normalization parameter for each model. We set external cAMP input concentrations in each model to the same values as in Figure 5C. A gray shaded background behind the plots highlights experimental data. A green shaded background indicates models that reproduce the experimental observations.

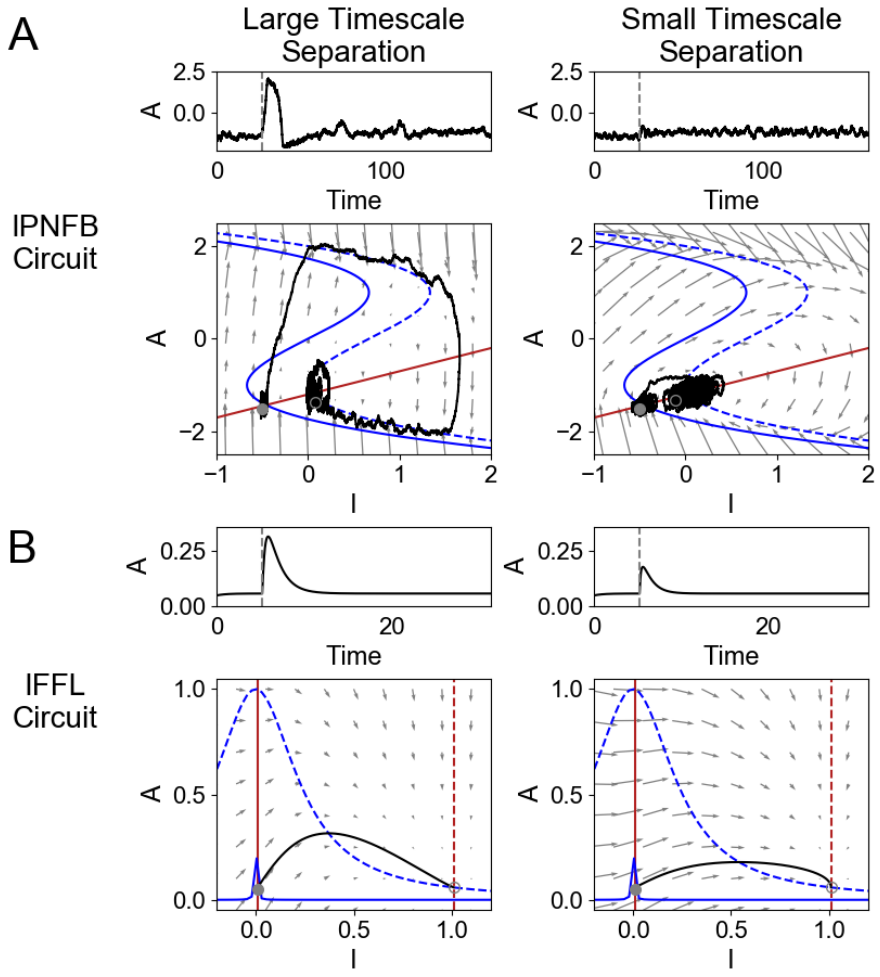


**FIG. S5: The effect of decreasing timescale separation on the single-cell response to step inputs.** Activator (approximating internal cAMP level) response traces and nullcline analysis with different timescale separation parameters in the IPNFB circuit **(A)** and the IFFL circuit **(B)**. Left and right panels show model activator responses for large (IPNFB $\varepsilon = 0.1$, IFFL $\tau = 1.5$) and small (IPNFB $\varepsilon = 1$, IFFL $\tau = 0.5$) values of the time scale separation parameter between the activator (A) and inhibitor (I) species, respectively. In simulations, we applied one unit of external cAMP at the gray dashed line. Solid black lines show how the activator trajectory in the phase plane. Solid gray dots and gray circles mark the initial and end states of the systems. Solid and dashed blue lines are activator nullclines before and after external cAMP input. Solid and dashed red lines are inhibitor nullclines before and after external cAMP input. Gray lines with arrows display vector fields of the systems. Both models are displayed in their native units from the original publications and given in *Model Equations and Parameters* [8, 17].
